# An Intrinsically Disordered Pathological Variant of the Prion Protein Y145Stop Transforms into Self-Templating Amyloids via Liquid-Liquid Phase Separation

**DOI:** 10.1101/2021.01.09.426049

**Authors:** Aishwarya Agarwal, Sandeep K. Rai, Anamika Avni, Samrat Mukhopadhyay

## Abstract

Biomolecular condensation via liquid-liquid phase separation of intrinsically disordered proteins/regions (IDPs/IDRs) along with other biomolecules is thought to govern critical cellular functions, whereas, aberrant phase transitions are associated with a range of deadly neurodegenerative diseases. Here we show, a naturally occurring pathological truncation variant of the prion protein (PrP) by a mutation of a tyrosine residue at 145 to a stop codon (Y145Stop) yielding a highly disordered N-terminal IDR that spontaneously phase-separates into liquid-like droplets. Phase separation of this N-terminal segment that is rich in positively charged and aromatic residues is promoted by the electrostatic screening and a multitude of other transient, intermolecular, noncovalent interactions. Single-droplet Raman measurements in conjunction with an array of bioinformatic, spectroscopic, microscopic, and mutagenesis studies revealed that the intrinsic disorder and dynamics are retained in the liquid-like condensates. Lower concentrations of RNA promote the phase transition of Y145Stop at low micromolar protein concentrations under physiological condition. Whereas, higher RNA to protein ratios inhibit condensation indicating the role of RNA in modulating the phase behavior of Y145Stop. Highly dynamic liquid-like droplets eventually transform into dynamically-arrested, ordered, β-rich, amyloid-like aggregates via liquid-to-solid transition upon aging. These amyloid-like aggregates formed via phase separation display the self-templating characteristic and are capable of recruiting and autocatalytically converting monomeric Y145Stop into amyloid fibrils. In contrast to this disease-associated intrinsically disordered Y145 truncated variant, the wild-type full-length PrP exhibited a much lower propensity for phase separation and liquid-to-solid maturation into amyloid-like aggregates hinting at a potentially crucial, chaperone-like, protecting role of the globular C-terminal domain that remains largely conserved in vertebrate evolution. Such an intriguing interplay in the modulation of the protein phase behavior will have much broader implications in cell physiology and disease.

## Introduction

A growing body of current research has revealed that in addition to canonical membrane-enclosed organelles, eukaryotic cells also contain organelles without delimiting membranes. These membrane-less organelles, also known as biomolecular condensates, are formed via liquid-liquid phase separation (LLPS) of proteins, nucleic acids, and other biomolecules.^1–16^ These mesoscopic intracellular compartments are dynamic, liquid-like, permeable, non-stoichiometric supramolecular assemblies that allow rapid exchange of components within the environment and offer unique spatiotemporal control of the macromolecular organization and cellular biochemistry. Intracellular phase transitions are governed by an intricate balance between the enthalpy and entropy of mixing and are primarily driven by intrinsically disordered proteins/region (IDPs/IDRs) containing prion-like low-complexity regions (LCRs) comprising a “stickers-and-spacers” sequence architecture.^1,17-28^ The sticker residues can promote weak, stereospecific, multivalent, transient intermolecular contacts involving electrostatic, hydrophobic, hydrogen bonding, dipole-dipole, π-π, and cation-π interactions. Whereas, the flexible spacer residues can dynamically control the making-and-breaking of noncovalent interactions giving rise to a liquid-like behavior of phase-separated biomolecular condensates. While liquid-like intracellular bodies are lined with a diverse array of cellular functions, liquid-to-gel and liquid-to-solid phase transitions are implicated in a range of debilitating neurodegenerative diseases.^29–39^ Therefore, there is an emerging consensus on the central role of biomolecular condensation and maturation promoting more persistent intermolecular contacts in aberrant pathological phase transitions.

The conformational conversion of the human prion protein (PrP) into a misfolded, β-rich, aggregated, self-replicating amyloid-like form is associated with a range of invariably fatal and transmissible neurodegenerative diseases classified under transmissible spongiform encephalopathies.^40,41^ This class of diseases is either caused by a spontaneous prion misfolding leading to sporadic Creutzfeldt-Jakob disease (CJD) or due to mutations in the *prnp* gene resulting in familial CJD, fatal familial insomnia, and Gerstmann-Sträussler-Scheinker (GSS) syndrome.^40-42^ Cellular PrP is 253-amino acids protein consisting of an N-terminal signal peptide (residues 1-23) that is cleaved upon maturation, a positively charged unstructured N-terminal tail (amino acids 23-120), a globular C-terminal domain (amino acids 121-231), and a GPI-anchor signal (residues 231-253) (Figure 1A,B).^40-43^ The N-terminal unstructured domain can be classified as an IDR that comprises two lysine clusters (residues 23-30 and 100-110), five glycine-rich octapeptide repeats (PHGGGWGQ), and a hydrophobic segment (113-135). In contrast, the C-terminal domain is highly structured and consists of three α-helices (amino acids 144-154, 175-193, and 200-219) and two short antiparallel β-strands (residues 128-131 and 161-164) (Figure 1B). *In vitro* prion conversions and amplifications from recombinantly expressed PrP recapitulate several key structural and biochemical characteristics of pathogenic prion deposits.^44–46^ An unusual pathological mutation of the tyrosine at residue 145 to a stop codon (PrP-Y145Stop) results in a C-terminally truncated intrinsically disordered variant of the protein that exhibits GSS-like phenotype.^47–49^ Under normal conditions, this fragment is unstable and is rapidly degraded in the cell, however, under stress conditions, Y145Stop accumulates in the endoplasmic reticulum, Golgi, and nucleus. This truncation mutation results in pathological amyloid deposits in the brain.^50,51^ In this work, we show that intrinsically disordered Y145Stop spontaneously phase-separates into highly dynamic liquid-like droplets that undergo a time-dependent maturation into ordered, solid-like, self-templating amyloid aggregates via a liquid-to-solid phase transition. The prion phase behavior can be modulated by RNA that tunes the material property of these condensates. The LLPS-mediated conformational switch of Y145Stop provides a mechanistic underpinning of effective nucleation and transition into self-replicating pernicious amyloid-like species.

**Figure 1.**
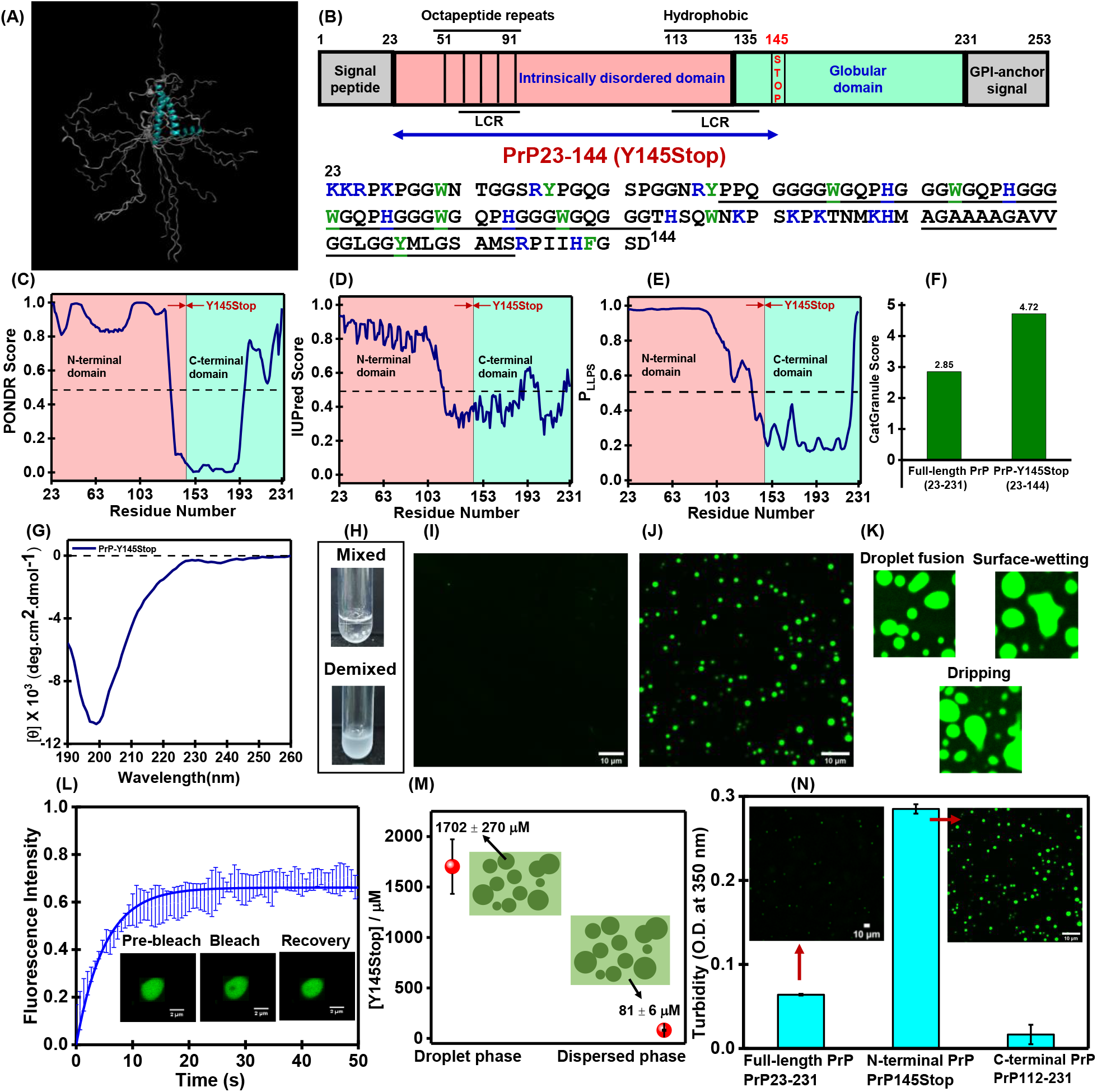
Phase separation of PrP-145Stop. (A) Overlay of 20 conformations obtained from the NMR structure of human PrP90-231 (PDB ID: 2LSB)^43^ generated using PyMOL (Schrödinger, LLC, New York) showing conformational heterogeneity at the N-terminal domain. (B) Schematic representation of PrP indicating all the segments and domains (Y145Stop is also highlighted). The amino acid sequence of Y145Stop from residues 23 to 144 showing positively charged residues (blue) and aromatic residues (green). Two low complexity regions (LCRs) predicted by SMART (Simple Modular Architecture Research Tool)^56^ are underlined. The disorder predictions tools PONDR^52^ (C) and IUPred^53^ (D) indicate highly unstructured N-terminal segment of PrP. A boundary between disordered and ordered regions is shown at residues 145 (this boundary is conventionally shown at residue 121). Prediction of LLPS propensity using FuzPred^54^ (E) and catGRANULE^55^ (F). The LLPS probability (P_LLPS_) of Y145Stop is 0.997 obtained from FuzPred. (G) A CD spectrum of 145Stop showing a random-coil conformation. (H) Spontaneous LLPS of Y145Stop (100 µM, pH 7.5, 37 °C) from a homogeneous mixed phase into a demixed phase upon the addition of salt (350 mM NaCl). Confocal fluorescence images of homogeneous phase (I) and droplets (J) performed using Alexa488-labeled Y145Stop single-Cys mutant at residue 31. (K) Droplet fusions, surface-wetting, and dripping. (L) FRAP kinetics of multiple droplets (using 10% Alexa488-labeled protein; n = 5). The solid line represents the fitted curve (See Supporting Info for details). (M) Concentration estimation of the Y145Stop droplet (dense) phase and dispersed (light) phase using centrifugation assays. Data are represented as mean ± SEM of independent measurements (n = 5). (N) Comparison of phase separation ability of Y145Stop, full-length PrP, and the C-terminal segment using turbidity and confocal microscopy. For 145Stop, the image shown in Figure 1J is included here for comparison.

## Results

### PrP Y145Stop undergoes liquid-liquid phase-separation *in vitro*

In order to examine the presence of intrinsic disorder and LCRs in Y145Stop, we first used well-known bioinformatic prediction tools^52-56^ which together revealed the presence of two LCRs (residues 50-94 and 113-135) and the highly disordered nature of Y145Stop. (Figure 1B-F). This was further verified experimentally by monitoring circular dichroism (CD) of recombinantly expressed human prion protein Y145Stop (Figure 1G). Next, we used phase separation prediction tools such as FuzPred^54^ and catGRANULE ^55^ that revealed a high LLPS propensity of intrinsically disordered Y145Stop (Figure 1E,F). To experimentally verify the LLPS propensity of PrP Y145Stop, we began by characterizing its phase behavior *in vitro* using turbidity measurements and confocal microscopy. Under cytosolic pH (pH 7.5) at 37 °C Y145Stop even at a high protein concentration (100 μM) remained clear and dispersed. Upon addition of increasing amounts of salt, the turbidity of the protein solution rose immediately indicating phase separation under cytosolic conditions (Figure 1H). We next created a single-Cys mutant at residue position 31 of Y145Stop in order to label the protein using a bright, photostable, thiol-active fluorescent dye, AlexaFluor488 maleimide. Y145Stop mixed with fluorescently-labeled protein (1%) was used for confocal imaging of droplets (Figure 1I,J). These fluorescent droplets undergo fusion, surface-wetting, dripping, and exhibit fast fluorescence recovery after photobleaching (FRAP) highlighting their liquid-like nature (Figure 1K,L). The concentration of protein inside droplets was estimated to be ∼ 2 mM, which is ∼ 25 times denser compared to the dispersed phase (∼ 80 µM) that corresponds to the critical saturation concentration, C_sat_ (Figure 1M). Fluorescence anisotropy measurements revealed no significant hindrance in the rotational relaxation of the polypeptide chain upon LLPS indicating a mobile interior within the droplets (data not shown). To mimic the cellular environment, we also studied phase separation of Y145Stop in the presence a of molecular crowding agent such as polyethylene glycol (PEG). In the presence of 10% PEG, Y145Stop undergoes LLPS at a much lower protein concentration and at a physiological salt concentration (150 mM) (Figure S1A).

Taken together, these results demonstrate that the N-terminal IDR of PrP, Y145Stop, under the near-physiological condition, undergoes condensation into liquid-like droplets with a protein-rich, dense, yet highly mobile, environment within the droplets. The net charge of this domain is + 10.4 at pH 7.5, and therefore, intermolecular repulsions between highly charged polypeptide chains prevent phase separation at low ionic strength. At higher salt concentrations, due to the effective charge-screening, polypeptide chains can commingle via weak, noncovalent, inter-chain interactions facilitating LLPS of this N-terminal IDR. Next, we asked what is the role of the C-terminal globular domain in modulating the prion phase behavior. PrP comprising both N- and C-terminal domains undergoes phase separation.^57,58^ However, under our condition, the phase separation propensity of the full-length PrP was much lower than the N-terminal IDR, Y145Stop (Figure 1N). This observation is consistent with the predictions (Figure 1E,F). We then created an N-terminally truncated form of the protein, PrP (112-231) which contains the entire C-terminal globular domain and a much shorter N-terminal tail comprising one of the two LCRs. As expected, this construct is well-folded into a helical structure, as evident by CD (Figure S1B). PrP (112-231) did not phase separate even after prolonged incubation under our conditions in the presence or absence of salt. Our experimental data corroborate the results obtained from the prediction tools and suggest that the C-terminal globular domain alone is not prone to LLPS. In fact, the presence of this globular domain in full-length PrP diminishes the LLPS propensity. Together these observations indicated that the N-terminal IDR is the principal driver of PrP phase separation, whereas, the C-terminal domain acts as an LLPS-inhibition domain indicating its chaperone-like role. Next, we set out to elucidate the key molecular determinants of phase separation of Y145Stop.

### Molecular drivers of phase separation of PrP Y145Stop

#### Electrostatic screening

As stated above, Y145Stop carries a high net positive charge (+10.4) at pH 7.5. Interchain repulsions prohibit intermolecular interactions that are essential for LLPS. In order to screen the high positive charge and to allow an effective interchain association, a threshold salt concentration is required above which it should phase separate via homotypic multivalent interactions. To unveil the role of electrostatic screening, we constructed the phase diagram as a function of salt concentration. Y145Stop phase separates above a critical threshold salt concentration (> 300 mM, pH 7.5) and remains phase-separated throughout the entire NaCl concentration range (Figures 2A,B, S2A). At a high salt concentration (750 mM NaCl), phase separation was observed at a much lower protein concentration (Figure 2B). In order to further support the role of electrostatic screening, we altered the charge-state of the protein by varying the solution pH at a fixed salt concentration. Both turbidity measurements and confocal imaging at different pH (pH 5.8, 7.5, and 8.8) at a fixed ionic strength showed an enhanced phase separation with an increase in the pH (Figure S2B,C). The protein at a mildly acidic pH (pH 5.8) carries a higher net positive charge (∼ +15), and therefore, requires a greater screening effect for phase separation compared to the protein at a mildly alkaline pH (pH 8.8) containing a lower net positive charge (∼ +8.5). These observations indicated that electrostatic screening is critical for phase separation of Y145Stop. An increase in the salt concentration decreases the Debye length allowing the screening of electrostatic repulsions between the positively charged residues resulting in the phase transition.^59^ Therefore, phase separation of Y145Stop is akin to ‘salting out’ like other phase separating proteins (TDP-43, hnRNPA1, and elastins) and is in contrast to the reentrant phase behavior in which LLPS is observed both at low salt and at high salt regime but not at intermediate salt concentrations.^60–62^ Since at high salt concentrations, the electrostatic interactions are negligible but the hydrophobic effect can be predominant,^63,64^ we next set out to characterize the role of hydrophobic interactions in driving phase separation of Y145Stop.

**Figure 2.**
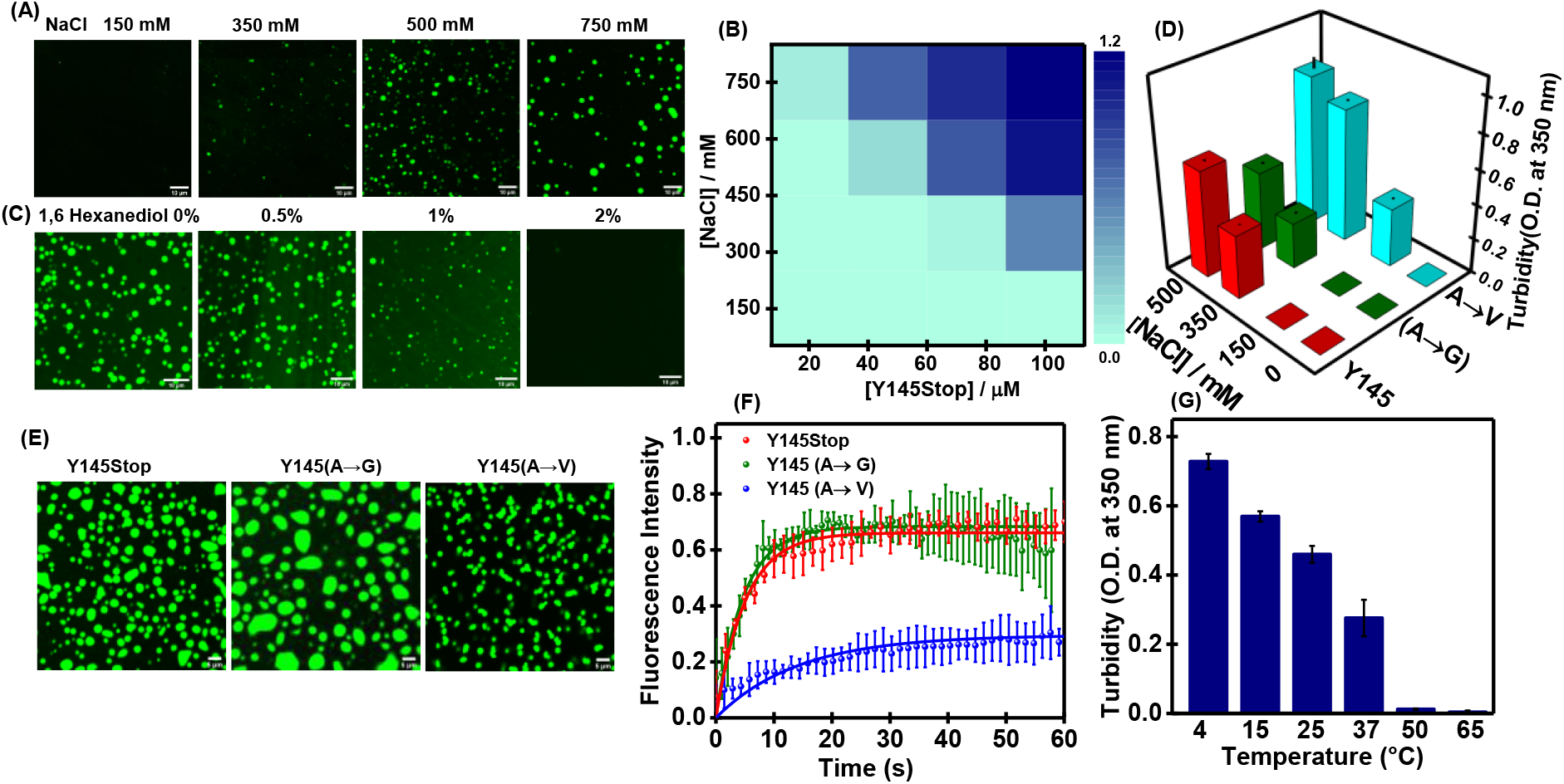
Molecular determinants of phase separation. (A) Confocal images of fluorescently labeled Y145Stop (100 µM, pH 7.5, 37 °C) as a function increasing NaCl concentrations. (B) A phase diagram at different protein and salt concentrations constructed from mean turbidity values. (C) Confocal images of droplets in the presence of increasing amounts of 1,6 hexane diol (100 µM Y145Stop, 350 mM NaCl, pH 7.5, 37 °C). (D) Turbidity plot for Y145Stop, and variants A→G and A→V in presence of different concentrations of salt. (E) Confocal images of Y145Stop and variants A→G and A→V (100 µM protein, 350 mM NaCl, pH 7.5, 37 °C) (F) FRAP kinetics of multiple fluorescently labeled droplets of Y145Stop and variants A→G and Y145 A→V (n = 5). (G) Phase separation of Y145Stop as a function of temperature.

#### Hydrophobic interactions

In order to characterize the role of hydrophobic interactions in promoting phase separation, we first used 1,6-hexanediol, an aliphatic alcohol, that is known to disrupt weak hydrophobic interactions. Phase separation was inhibited by 2% (w/v) 1,6-hexanediol indicating an important role of hydrophobic interactions in LLPS of Y145Stop (Figure 2C, S3A). We also used ANS (8-anilino-1-naphthalenesulfonic acid) that is a well-known hydrophobic reporter. We observed a blue-shift and an increase in the ANS fluorescence intensity upon LLPS of Y145Stop indicating the presence of local hydrophobic patches within the droplets (Figure S3B). A closer look at the amino acid sequence of Y145Stop revealed the presence of a hydrophobic stretch in the second LCR (residues 112-121) that is rich in alanine and valine (Figure 1B). We further evaluated the role of hydrophobicity by performing site-directed mutagenesis at the hydrophobic stretch. We mutated residues by substituting three alanines to valines to increase the hydrophobicity and alanines to glycines to decrease the hydrophobicity. These mutations did not alter the structural attributes of Y145Stop as verified using CD measurements (Figure S3C). Glycine is polar and is a well-known spacer that does not appear to alter the saturation concentration.^65^ As expected, the substitution of alanines to glycines (Y145StopA→G) afforded droplets with a bit enhanced fluidity and behaved similarly to the wild-type Y145Stop (Figure 2D,E). However, the mutations of alanines to valines (Y145StopA→V) considerably enhanced the LLPS propensity and resulted in more solid-like and aggregation-prone droplets as revealed by irregular shape in confocal microscopy associated with much lower FRAP recovery (Figure 2F). Thus, the mutations within the hydrophobic domain alter the phase behavior and simultaneously modulate the material properties of the assemblies. Together, these results indicate the role of hydrophobicity in driving the phase separation of Y145Stop. If the hydrophobic effect is a primary molecular driver of the phase transition, we would expect to observe higher phase separation propensity at higher temperatures due to the increase in the entropic contributions. Therefore, our next goal was to examine the thermo-responsive phase behavior of Y145Stop.

#### Thermo-responsive phase behavior

The interplay of molecular drivers of phase separation is critically dependent on the amino acid composition.^20,65-68^ Based on the amino acid composition, IDPs exhibit the upper critical solution transition (UCST) and the lower critical solution temperature (LCST).^67^ Polar LCRs interspersed by aromatic and charged residues display the UCST behavior, whereas, hydrophobic/aromatic-rich sequences follow the LCST behavior since the hydrophobic effect being largely entropic is more pronounced at a higher temperature. In order to characterize the temperature-dependent phase transition of Y145Stop, we performed the turbidity assay as a function of temperature ranging from 4 °C to 65 °C. Interestingly, we observed an increase in the phase separation at lower temperatures indicating a UCST behavior (Figure 2G). These droplets exhibited temperature-dependent reversibility in the phase transition (Figure S4A). An LCST behavior is expected for LLPS that is primarily driven by hydrophobic interactions. The amino acid sequence of Y145Stop contains polar LCRs enriched in glycine, serine, glutamine, and asparagine, along with aromatic amino acids (tryptophan, tyrosine, and phenylalanine) and positively charged residues (lysine and arginine). The anomalous thermo-responsive characteristics plausibly point towards the dual behavior of aromatic amino acids central to both LCST and UCST transitions. These aromatic residues can participate in intermolecular π-π stacking and cation-π interactions to promote LLPS.^65^ Our phase separation assays using free arginine showed that LLPS of Y145Stop is abolished at higher concentrations of arginine presumably due to strong interactions of free arginine with the aromatic sidechains that can potentially compete with intermolecular cation-π interactions (Figure S4B,C). These results indicate the participation of aromatic residues in promoting phase separation of Y145Stop. Taken together, this set of studies highlight the presence of promiscuous interactions within the condensed environment of liquid-like droplets. The electrostatic screening allows the key noncovalent forces to promote transient, multivalent, interchain interactions between highly flexible conformers driving the liquid phase condensation of Y145Stop. Our next goal was to discern the key structural characteristics of polypeptide chains within the liquid droplets.

### Vibrational Raman spectroscopy captures the key conformational signatures

In order to elucidate the protein conformational states, we employed vibrational Raman spectroscopy that provides a wealth of molecular information about the polypeptide backbones and sidechains.^69,70^ Additionally, a laser Raman assembly equipped with a microscope allows us to focus the laser beam into the protein-rich, dense, condensed phase of individual droplets to capture Raman scattering bands associated with a multitude of molecular vibrational modes from protein-rich single droplets. The Raman spectra of dispersed and phase-separated states of Y145Stop clearly showed characteristic bands corresponding to backbone amide I, amide III, Trp, Tyr, Phe, and other vibrational modes (Figure 3A). Amide I (1630-1700 cm^−1^) originates primarily due to the C=O stretching of the polypeptide backbone, whereas, amide III (1230-1300 cm^−1^) corresponds to a combination of N-H bending and C-N stretching modes.^69,70^ These amide vibrational bands are often used to identify the secondary structural elements in proteins. Broad amide I bands for both dispersed and demixed phases exhibited highly disordered conformations. A closer inspection of amide I revealed an increase in the FWHM (full-width at half maximum) from the dispersed phase (∼ 55 cm^−1^) to the droplet phase (∼ 63 cm^−1^) indicating a higher conformational heterogeneity in the condensed phase (Figure 3B). Next, we determined the intensity ratio of the tyrosine Fermi doublet (I_850_/I_830_) that serves as an indicator of the hydrogen bonding strength between the phenolic hydroxyl group of Tyr and the neighboring water molecules in the vicinity. This ratio exhibited a small decrease from ∼ 2.75 to ∼ 2.67 upon phase separation indicating a small but measurable drop in the solvent accessibility in the liquid-like condensed phase. The intensity ratio for the tryptophan Fermi doublet (I_1360_/I_1340_) (hydrophobicity) and a band at 880 cm^−1^ (hydrogen bonding strength between water and N-H of the indole ring) did not exhibit any measurable changes upon the liquid phase condensation. We were able to recapitulate all these vibrational signatures in our single-droplet Raman studies. Taken together, these results demonstrate that Y145Stop retains its intrinsic disorder with a slight increase in the structural heterogeneity within the demixed liquid droplets.

**Figure 3.**
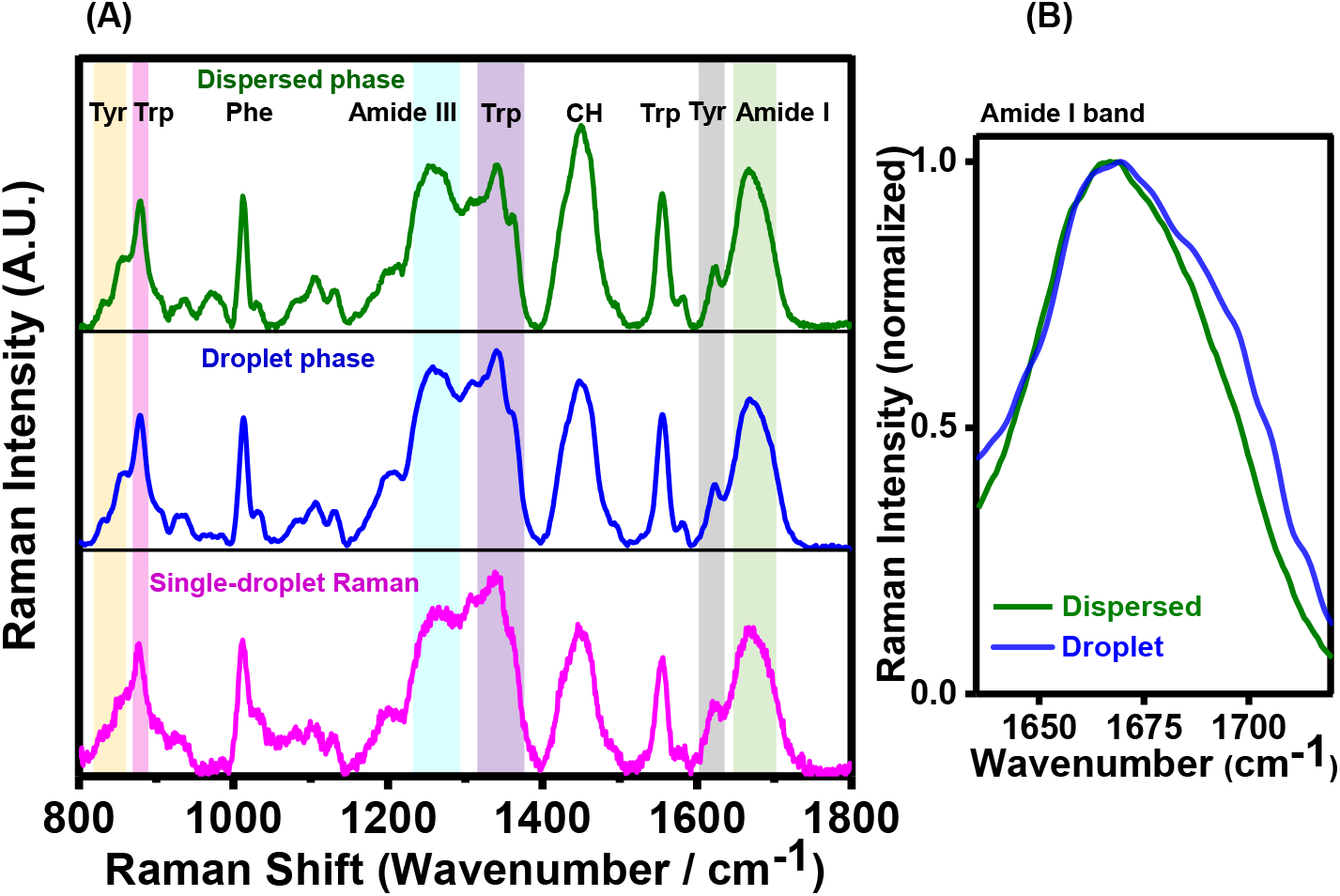
Vibrational Raman spectroscopy of Y145Stop. (A) Bulk Raman spectra of dispersed (olive) and droplet phases (blue) and single-droplet Raman (magenta). Prominent vibrational signatures of amide backbones and sidechains are shaded and labeled. (B) The amide I regions are shown for dispersed and condensed phases.

### RNA regulates the phase behavior of Y145Stop

The N-terminal IDR, Y145Stop, is broadly classified under the umbrella of RNA-binding proteins due to the presence of two highly conserved polybasic regions containing lysine clusters that have charge complementarity to nucleic acids.^71,72^ We next examined the role of RNA in Y145Stop phase separation. The addition of crude t-RNA as low as 100 ng resulted in phase separation as confirmed using turbidity assays and confocal microscopy. The presence of RNA drastically lowered the saturation concentration of the protein to as low as 5 μM in the presence of 150 mM NaCl (Figure 4A,B). The droplets were initially spherical and rapidly transformed into a gel-like state comprising non-spherical condensates. We next performed FRAP experiments to investigate the protein diffusivity within these RNA-protein condensates. We observed much lower and incomplete recovery of these RNA-induced droplets (∼ 20%) as compared to salt-induced droplets (∼ 70%) indicating a more hardened material property of RNA-Y145Stop condensates (Figure 4C). In order to rule out the possibility of specificity of RNA in the observed phase behavior of Y145Stop, we also performed phase separation assays in the presence of other RNAs such as poly-U RNA and yeast total RNA. Our results showed a similar ability of these RNAs in modulating the phase behavior of Y145Stop (Figure S5). Next, to establish if the RNA-induced LLPS resulted in a more viscous, gel-like interior, we recovered apparent diffusion coefficients of the protein within the droplets (See Supporting Info).^34^ In the absence of RNA, the FRAP recovery was on a faster timescale with a t_1/2_ of 3.48 s that yielded an approximated diffusion coefficient of 0.027 µm^2^s^−1^. The recovery kinetics in the presence of RNA was much slower (t_1/2_ = 6.96 s) indicating a considerable drop in the apparent diffusion coefficient (0.013 µm^2^s^−1^). Therefore, the RNA-induced droplets are associated with lower protein diffusivity and enhanced internal viscosity. Next, we studied the effect of the RNA concentration on the phase separation of Y145Stop. We found that the phase separation ability of the protein is highly enhanced at a low RNA concentration, whereas, at higher concentrations of RNA, no phase separation was observed (Figure 4D,E). Protein solutions with high RNA concentrations remained dispersed and perfectly mixed even upon longer incubation. The interaction of poly-anions such as RNAs with the positively charged N-terminal domain possibly results in the charge inversion that prevents phase separation. Such an RNA stoichiometry-dependent promotion and inhibition of phase transition has been previously observed for FUS, TDP-43, LAF-1, and so forth.^73–78^ Together these results suggest that RNA buffers the phase behavior of Y145Stop and tune the material property of the condensates.

**Figure 4.**
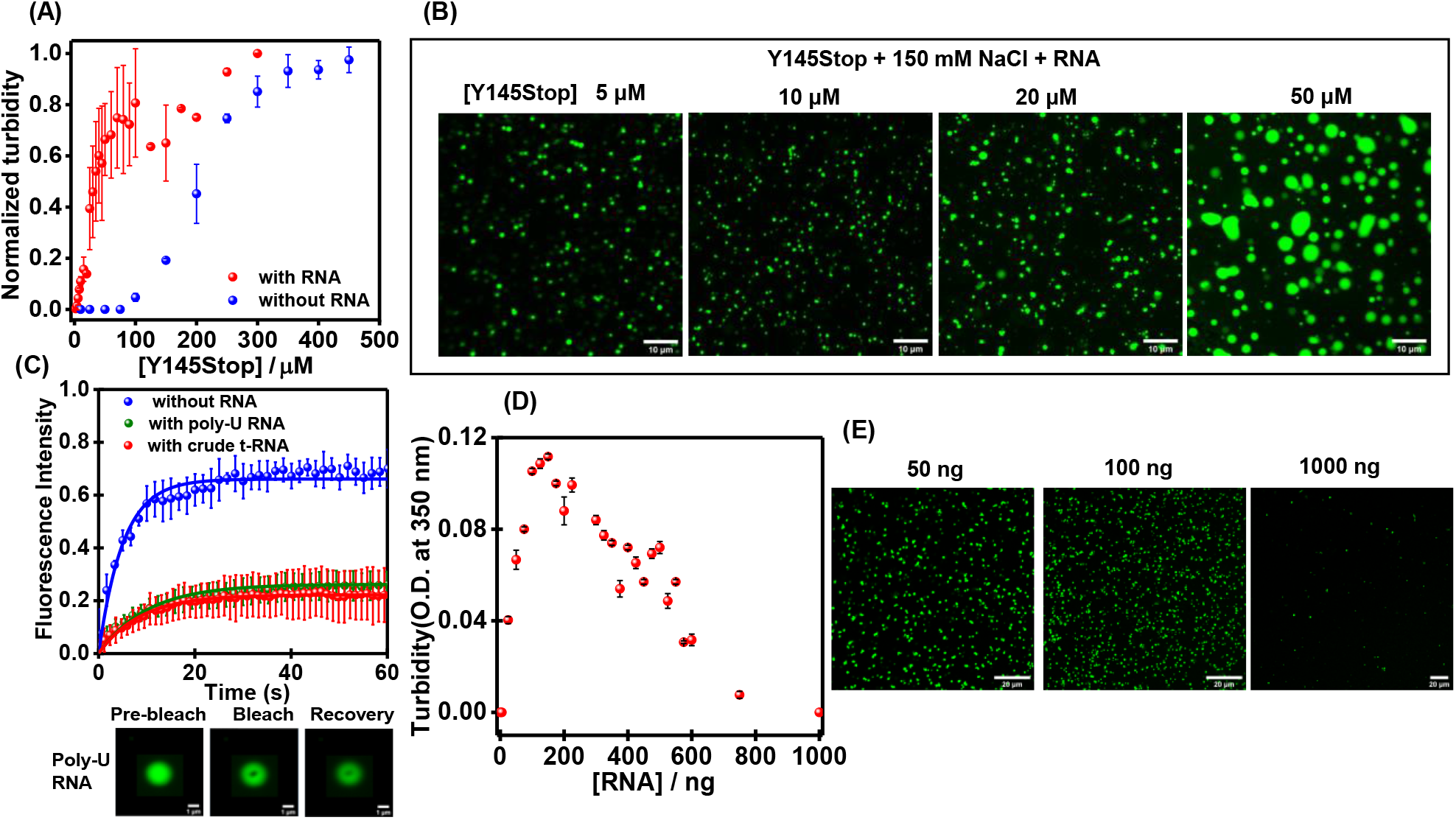
Modulation of Y145Stop phase transitions by RNA. RNA lowers the Y145Stop concentration required for phase separation from ∼ 80 µM to 5 µM of protein at 150 mM NaCl and pH 7.5 as observed by the turbidity assay (A) and confocal microscopy (B). (C) FRAP kinetics of multiple Y145Stop droplets (10% Alexa488 labeled Y145Stop, n=5) in the absence and the presence of RNA (crude t-RNA and poly-U RNA). The solid lines represent the fitted curves (see SI). The fluorescence images of droplets during FRAP measurements are also shown. (D & E) Low RNA/protein ratios promote phase separation and high ratio inhibit phase separation at a fixed (10 µM) Y145Stop concentration as observed by the RNA-concentration-dependent turbidity assays and confocal microscopy (poly-U RNA).

### Phase separation of Y145Stop promotes the formation of amyloid-like aggregates

In order to monitor the maturation of Y145Stop condensates, we monitored the aging morphology of liquid droplets under a confocal microscope (Figure 5A). The liquid-like characteristics of these droplets were retained till ∼ 5 h after which these droplets irreversibly transformed into gel-like and/or solid-like fibrous morphologies. We observed ‘sea urchin’-like structures that were previously observed for FUS aggregates formed via LLPS.^79^ These results indicate that the droplets might act as the centers for nucleation and growth of fibers (Figure 5A). The solid-like nature of these aggregates was also confirmed using FRAP which showed no recovery (Figure 5B,C). These aggregates formed via the aging of liquid-like condensates are thioflavin-T (ThT) positive indicating a transition into amyloid-like aggregates (Figure S6A). Our vibrational Raman spectroscopic investigation showed a sharp amide I band at 1675 cm^−1^ that is a hallmark of hydrogen-bonded cross-β architecture in amyloid fibrils (Figure 5D).^70^ The FWHM of amide I considerably narrowed down from liquid droplets (∼ 63 cm^−1^) to aggregates (∼ 44 cm^−1^) indicating a more ordered and less heterogeneous conformational state. The ratio of the tyrosine fermi doublet (I_850_/I_830_) decreased considerably upon liquid-to-solid transition from droplets (∼ 2.67) to aggregates (∼ 2.16) suggesting a buried environment of Tyr residues with a lower propensity to form hydrogen bonds with water. Additionally, the polarity around Trp also exhibited a small decrease as evident by a slight blue shift of the band around 880 cm^−1^ indicating a lower hydrogen bonding ability of N-H of the indole ring with water within the aggregates. Together these results suggested conformational sequestration of disordered polypeptide chains into a well-ordered amyloid architecture formed via liquid phase condensation and liquid-to-solid maturation.^80^ This LLPS-mediated liquid-to-solid transition is also characterized by the time-dependent evolution of intrinsic blue fluorescence (Figure 5E) that arises due to the intermolecular charge-transfer and electron delocalization via extensively hydrogen-bonded amide backbones within the phase-separated assemblies.^81^ Our atomic force microscopy (AFM) imaging revealed a fibrillar nanoscale morphology along with some amorphous aggregates (Figure 5F). Such amyloid fibrils with a diameter ∼ 10 nm were also observed upon the phase transition and aging in the presence of RNA at a much lower protein concentration (Figure S6B). We also examined the aggregates formed via the aging of liquid droplets of the full-length PrP under the same condition. These PrP aggregates exhibited much lower ThT intensity compared to Y145Stop indicating the C-terminal folded domain lowers the aggregation propensity (Figure S6C). This observation hints at a protective, chaperone-like role of the C-terminal globular domain in the absence of which the protein undergoes an aberrant phase transition into amyloid-like aggregates. We next asked whether these amyloid aggregates formed via phase separation of Y145Stop can exhibit a self-templating autocatalytic behavior that is a key characteristic of prion-like mechanisms.

**Figure 5.**
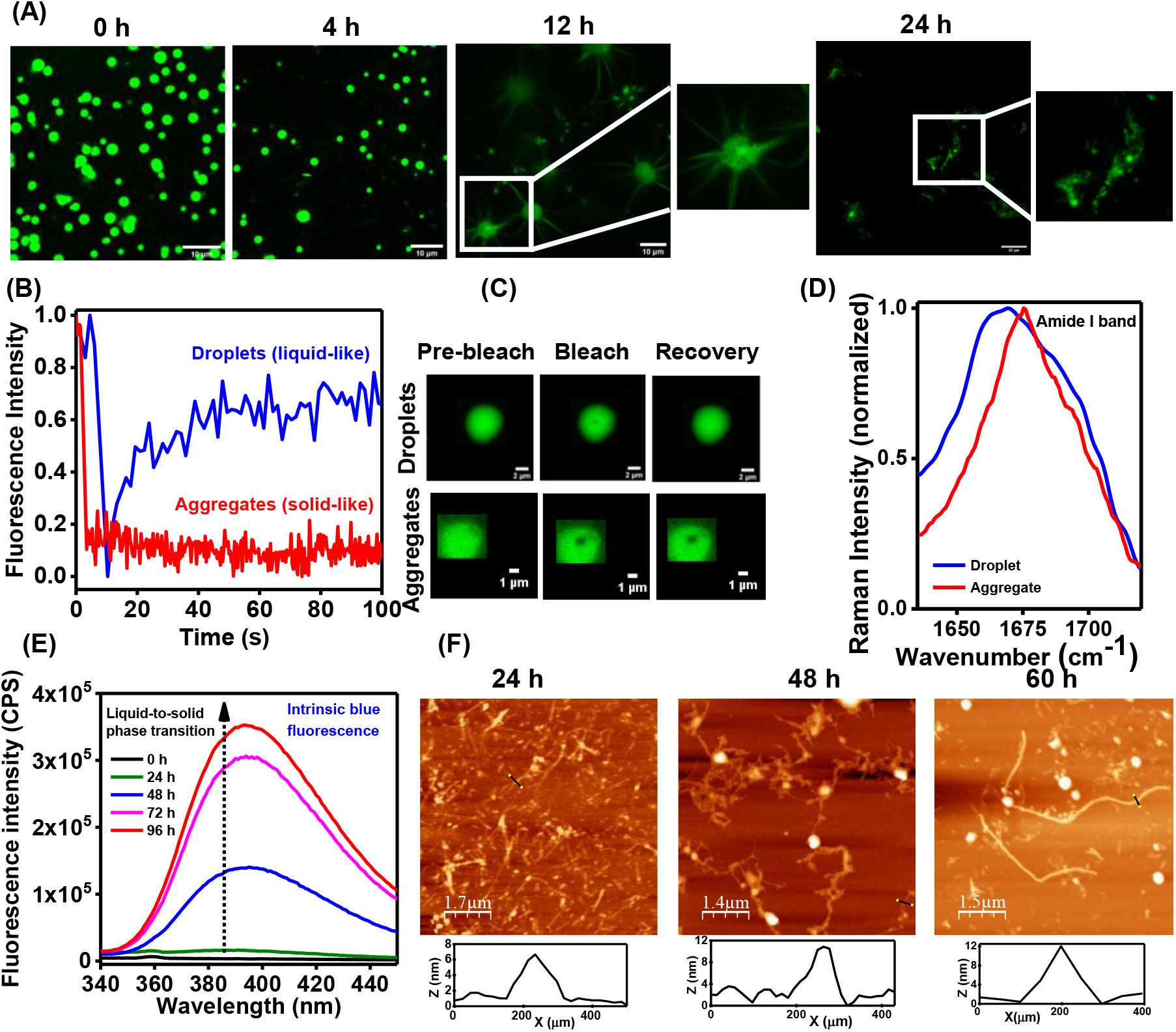
Liquid-to-solid phase transition of Y145Stop (A) Confocal images of fluorescently labeled Y145Stop showing morphological transitions as a function of time. (B) FRAP kinetics of solid-like aggregates showing no recovery (a representative FRAP kinetics of liquid-like state is also shown for comparison). (C) The fluorescence images during FRAP measurements. (D) Amide I vibrational Raman band showing a narrow peak at 1675 cm^−1^ indicating an amyloid-like cross-β architecture. (E) Time-evolution of the intrinsic blue fluorescence during maturation. (F) Time-dependent AFM images of aggregates formed via LLPS showing the presence of typical amyloid fibrils with heights ranging from 8-12 nm.

### Aggregates formed via phase transition display a self-perpetuating autocatalytic behavior

In order to probe the autocatalytic conversion, we first carried out a *de novo* aggregation reaction with Y145Stop that is known to proceed via a typical nucleation-dependent polymerization mechanism.^47^ The *de novo* aggregation kinetics exhibited a lag time ∼ 6 h. Next, we carried out these kinetic experiments in the presence of increasing amounts of preformed seeds obtained from the salt-induced LLPS-mediated phase transition and aging for 24 h. Upon addition of these seeds (0.5% −10%), the lag phase shortened in a dose-dependent manner (Figure 6A). By using 10% seeds in the reaction mixture, we were able to nearly bypass the lag phase ensuing a quasi-pseudo first-order kinetics. After establishing that the aggregates formed via phase separation are capable of recruiting and inducing a self-perpetuating conformational conversion, we asked which key species during liquid-to-solid maturation exhibit the most potent self-templating characteristics. In order to answer this question, we performed aggregation assays using the preformed seeds generated by the phase separation process followed by different aging times. Liquid droplets formed immediately upon LLPS exhibited much weaker seeding ability compared to the aged samples that were prepared by maturation into β-rich aggregates via liquid-to-solid transitions (Figure 6B). This observation suggests that amyloid conformers formed via the disorder-to-order phase transition are the key species in the autocatalytic conversion. The aggregation reaction seeded with aged solid-like species yielded typical amyloid fibrils reminiscent of nanoscale morphologies identical to fibrils formed via an unseeded *de novo* aggregation process (Figure 6C). Together these findings reveal that aberrant phase transitions of Y145Stop can produce self-replicating amyloid species recapitulating the *bona fide* prion-like behavior.

**Figure 6.**
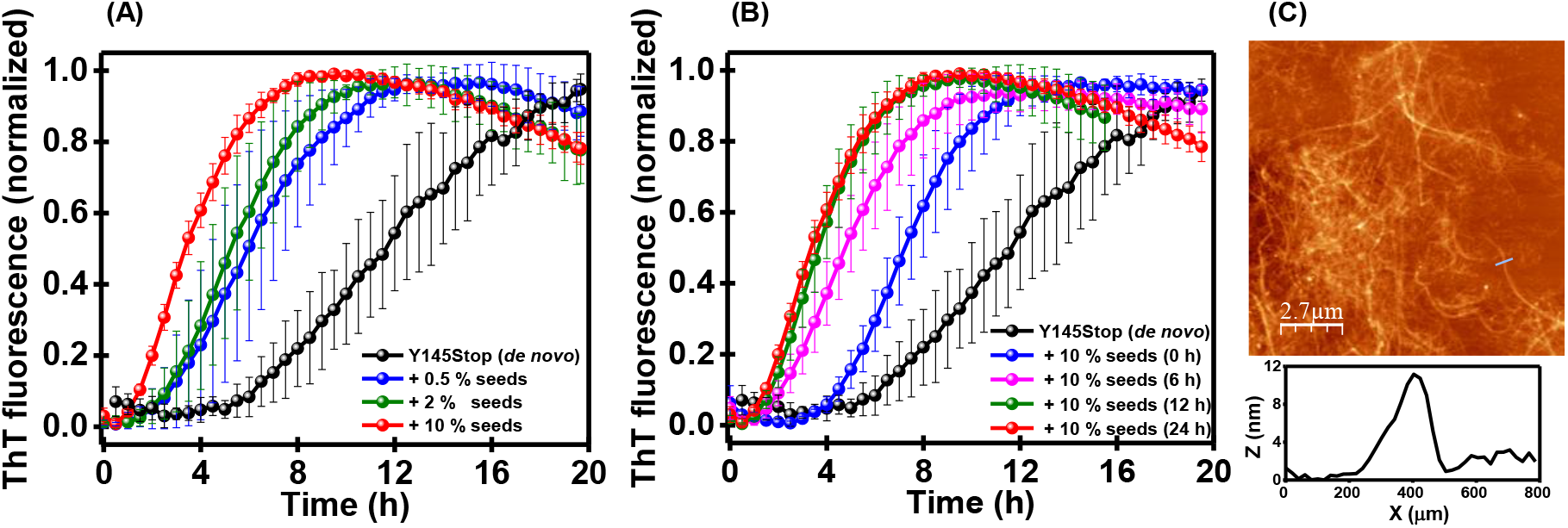
Seeding behavior of aggregates formed via LLPS. (A) The shortening of lag phase of *de novo* aggregation of Y145Stop (100 μM) by the addition of increasing amounts of amyloid seeds obtained from phase separation of Y145Stop. (B) The seeding efficiency monitored as a function of aging of phase separated Y145Stop indicating the most effective seeding after phase maturation. (C) The AFM image of Y145Stop fibrils with the height profiles obtained after seeded assembly.

## Discussion

In this work, we showed that the intrinsically disordered PrP-145Stop containing oligopeptide repeats and LCRs undergoes an instantaneous phase separation at near-neutral pH concordant with the bioinformatic predictions. These phase-separated condensates formed immediately after inducing LLPS exhibit liquid-like characteristics having a highly mobile internal organization. The electrostatic screening of this highly positively charged N-terminal fragment at a higher salt concentration allowed the polypeptide chains to interact via weak and transient intermolecular contacts driving a reversible, thermo-responsive phase transition. The hydrophobic effect along with other noncovalent interactions serve as the physical crosslinks essential for phase transitions. Single-droplet Raman measurements revealed that the conformational disorder and structural heterogeneity are retained within the liquid-like droplets. Phase separation is further promoted by the addition of a molecular crowding agent at much lower protein and salt concentrations mimicking the cytosolic milieu. Upon aging, the liquid-like droplets gradually transform into irreversible gel-like and solid-like ordered fibrillar aggregates that exhibit amyloid-like characteristics. These amyloids formed via phase separation of Y145Stop display the self-templating autocatalytic behavior that is a key characteristic of the prion-like propagation mechanism. Full-length PrP containing both N-terminal IDR and C-terminal globular domain undergoes phase separation as observed before,^57,82,83^ albeit with a much lower propensity compared to Y145Stop under the same condition. This observation also corroborates the bioinformatic prediction. Additionally, the propensity for the liquid-to-solid transition into amyloid-like form is also much lower for the full-length protein than the N-terminal fragment. Taken together, our results showcase the crucial role of the Y145Stop variant comprising the N-terminal IDR in driving phase separation and aberrant phase transitions.

Our findings reveal that RNA critically modulates the phase behavior of Y145Stop that contains putative RNA binding sites. Low RNA/protein ratios promote phase separation at a much lower Y145Stop concentration (5 μM) under the near-physiological condition (pH 7.5, 150 mM NaCl). RNA being a polyanion has a more pronounced influence in electrostatic neutralization of highly positively charged IDR than screening by salt ions in the absence of RNA. Therefore, RNA binding allows the chains to interact via a multitude of noncovalent interactions resulting in a complex coacervation. At low RNA/protein ratios, the RNA-induced droplets are more gel-like and rapidly transform into solid-like aggregates devoid of internal fluidity. In contrast, at higher RNA/protein ratios, the phase separation ability of Y145Stop was abolished. These observations suggested the stoichiometry of RNA and protein is critical in the RNA-dependent phase transition of Y145Stop. Our findings also highlight a broader role of RNA in aberrant phase transition and amyloid formation of Y145Stop that is associated with the GSS-like phenotype. The truncated Y145Stop variant is normally degraded by the proteasomal machinery or acted upon by the molecular chaperones. However, an impairment in the protein quality control allows this variant to accumulate in the endoplasmic reticulum, Golgi, and nucleus. Y145Stop overexpressed in cells treated with proteasome inhibitors has been shown to accumulate in the nucleus.^51,84^ Its presence in the nucleus is analogous to the aggregates found in diseases with polyglutamine repeats expansion. The presence of Y145Stop aggregates in the nucleus is intriguing as it has been ascribed to a cryptic nuclear localization signal which is otherwise masked in the full-length PrP.^51^

In summary, our study illuminates an intriguing interplay of molecular determinants that promote and regulate the phase transition and maturation into ordered self-templating amyloids from a pathological truncation variant of the prion protein. We would like to note that the putative functions of PrP such as copper homeostasis in the brain involve the binding of Cu^2+^ with the oligopeptide repeats located at the N-terminal IDR.^85,86^ Additionally, other proposed functions involving the maintenance of synaptic plasticity, suppressing apoptosis, resistance to oxidative stress are also located at the N-terminal domain.^85,86^ Even though the prion functions are mainly localized at the N-terminal segment, the C-terminal folded domain remains highly conserved during the vertebrate evolution. We, therefore, posit that the C-terminal globular domain can potentially have a protective chaperone-like activity to maintain the solubility of the functional N-terminal domain preventing its aberrant phase transitions. Additionally, our results underscore the crucial role of phase-separated droplets as the reaction crucibles for efficient nucleation and effective sequestration into self-perpetuating amyloid conformers. These critical molecular events can have broad implications in biological phase transitions associated with physiology and disease.

## Acknowledgments

We thank IISER Mohali, Department of Science and Technology (Nano-Mission grant to S.M.), Department of Biotechnology (fellowship to A.A.), Council of Scientific and Industrial Research (fellowship to S.K.R.), Ministry of Education, Govt. of India (Centre of Excellence grant to S.M.) for financial support, Prof. Witold Surewicz (Case Western Reserve University) for kind gift of the DNA plasmid of full-length PrP, Dr. Mahak Sharma (IISER Mohali) for providing access to confocal microscope, Ms. Pallavi Joshi for preliminary studies, Dr. Shravan K. Mishra (IISER Mohali) for providing yeast total RNA, and Dr. Mily Bhattacharya (Thapar Institute) and the members of the Mukhopadhyay lab for critically reading this manuscript.

