## Supporting Information for "An Intrinsically Disordered Pathological Variant of the Prion Protein Y145Stop Transforms into Self-Templating Amyloids via Liquid-Liquid Phase Separation"

### Materials and Methods

#### Materials

Sodium phosphate monobasic dihydrate, sodium phosphate dibasic dihydrate, trizma base, 1,6-Hexanediol, poly(ethylene glycol) (PEG), L-arginine, L-lysine sodium salt, sodium hydroxide, sodium chloride, L-glutathione reduced, 2-mercaptoethanol, 8-anilino-1-naphthalenesulfonic acid (ANS), Thioflavin T, 1,4-dithiothreitol (DTT), Triton X-100, Thrombin from bovine plasma, Poly-U sodium salt, phenylmethylsulfonyl fluoride (PMSF), ethylenediaminetetraacetic acid (EDTA), were of highest purity grade, obtained from Sigma (St. Louis, MO, USA). Crude t-RNA was purchased from Sisco Research Laboratory. Urea and guanidinium hydrochloride were procured from Amresco. Ampicillin, chloramphenicol, and isopropyl-β-thiogalactopyranoside (IPTG) were purchased from Gold Biocom (USA). All the fluorescent probes used in this study namely, fluorescein-5-maleimide (F-5-M), AlexaFluor488-C5-maleimide, and AlexaFluor594-C5-maleimide, were obtained from Molecular Probes, Invitrogen. Ni-NTA resin was purchased from Qiagen. PD-10 column was obtained from GE Healthcare Life Sciences (USA). Amicon membrane filters for concentrating protein were procured from Merck Millipore. All the buffer solutions were prepared in Milli-Q water and filtered before use. The pH of each buffer solution was adjusted (± 0.01) at 25 ° C using a Metrohm 827 lab pH meter.

#### Methods

##### Bioinformatic analyses of prion protein using various prediction tools

PONDR[1] and IUPred2a[2] were used to predict the intrinsic disorder in the prion protein (PrP). PONDR-VLXT and IUPred2a files were generated using PONDR (http://www.pondr.com/) and IUPred2a (https://iupred2a.elte.hu/) respectively and were then plotted using Origin 2018. FuzPred[3] and catGRANULE[4] were used to predict the LLPS propensity in the prion protein. The data were generated using FuzPred (http://protdyn-fuzpred.org/) and catGRANULE (http://s.tartaglialab.com) and were plotted using Origin 2018. SMART (Simple Modular Architecture Research Tool) (http://smart.embl-heidelberg.de/) was used to identify the LCRs (low-complexity regions).[5]

**Site-directed mutagenesis**

Y145Stop mutation was created using human PrP (23-231) plasmid which was a kind gift from Prof. Witold K. Surewicz (Case Western Reserve University, USA). Single cysteine variant of Y145Stop (W31C), Y145Stop (A→G) (A113A115A117 to G113G115G117), Y145Stop(A→V) (A113A115A117 to V113V115V117) variants of Y145 stop mutants were created using QuickChange site-directed mutagenesis kit (Stratagene). The primers used for the respective mutations are listed in Table1. All the mutations were verified by sequencing and the protein constructs were characterized by Circular Dichroism.

**Recombinant protein expression and purification**

Proteins were expressed and purified using previously published protocols with modifications.[6,7] Briefly, cleavable His-tagged recombinant Y145Stop, Y145Stop (A→G), Y145Stop (A→V), PrP (112− 231), and PrP (23-231) cloned in the pRSET-B plasmid were transformed into *E. coli* strain BL21(DE3) pLysS. Bacterial cultures were grown in LB media at 37 ˚C, 220 rpm. At $O.D._{600}$ = 0.6, the culture was induced with 1 mM IPTG for 8 hours at 30 ˚C, 220 rpm. Cell pellets were harvested by centrifugation at 4 ˚C, 4000 rpm for 30 minutes. These pellets were stored at -80 ˚C for further use. Purification for His-tagged constructs was performed using Ni-NTA chromatography as described previously.[6] Y145Stop(A→V) was eluted with buffer containing 500 mM imidazole in 8 M Urea (8 M urea, 10 mM Tris, 100 mM sodium phosphate, pH 8.0). The cysteine mutant Y145Stop (W31C) was purified under a denaturing condition from inclusion bodies as follows. Briefly, the pellets were resuspended in lysis buffer (50 mM tris-HCl, 100 mM NaCl, 1 mM EDTA, 0.1 % Triton X -100, pH 8.0) and cell-lysis was done by sonication using a probe sonicator (5% amplitude, 15 seconds on and 10 seconds off pulses, for 25 minutes). The supernatant was removed by centrifugation and pellets were washed twice. The pellets were then dissolved in denaturation buffer and were incubated for ~ 2-3 h. The supernatant collected after centrifugation was loaded onto the pre-equilibrated Ni-NTA column. Protein was eluted with buffer containing 500 mM imidazole (8 M urea, 10 mM Tris-HCl, 100 mM sodium phosphate, pH 7.5). Histidine tail-fused protein was dialyzed against buffer: 20 mM sodium phosphate, 50 mM NaCl, pH 6.4. The N-terminal His-tag was removed by incubating the protein with 0.2 U/mL of the thrombin protease at 37 ˚C for 5 hours. After the completion of the cleavage reaction, thrombin was inactivated by adding 0.2 mM PMSF in the protein solution. Further, to separate the His-cleaved and uncleaved protein, the protein was loaded onto the Ni-NTA column and eluted with 20 mM imidazole buffer H (8 M urea, 10 mM Tris-HCl, 100 mM sodium phosphate, pH 8.0). Protein was concentrated using 3 kDa MWCO amicon membrane filters and buffer exchanged (20 mM phosphate buffer, pH, 7.5) using PD10 columns. The purity of the protein was confirmed by SDS-PAGE analysis. The concentration of the protein were estimated using $\varepsilon_{280\ nm}$ = 43,670 $M^{-1}cm^{-1}$ for Y145Stop, Y145Stop(A→G) and Y145Stop(A→V), $\varepsilon_{280\ nm}$ = 56,590 $M^{-1}cm^{-1}$ for PrP(23-231), $\varepsilon_{280\ nm}$ = 14,200 $M^{-1}cm^{-1}$ for PrP (112-231) . To avoid the freeze-thaw cycles, all the experiments were performed using freshly purified proteins.

**Fluorescence labeling**

For labeling purified cysteine mutant in denaturing buffer was mixed in molar ratio of 10:1 (F-5-M:Y145Stop[W31C]), 2:1 (AlexaFluor488-C5-maleimide:Y145Stop[W31C]), and 2:1 (AlexaFluor594-C5-maleimide:Y145Stop[W31C]). The reaction mixtures were stirred for 2-3 h in dark at room temperature. After completion of labeling reaction, the excess free dye was removed using a PD10 column. The concentration of the labeled protein was estimated using $\varepsilon_{495\ nm}$ = 68,000 $M^{-1}cm^{-1}$, for F-5-M, $\varepsilon_{493\ nm}$ = 72,000 $M^{-1}cm^{-1}$, for AlexaFluor488 C5-maleimide, and $\varepsilon_{588\ nm}$ = 96,000 $M^{-1}cm^{-1}$ for AlexaFluor488 C5-maleimide.

**Phase separation assays**

Phase separation was immediately induced by adding NaCl to the reaction mixture incubated at 37 ˚C. The turbidity of the phase-separated samples was estimated by taking absorbance at 350 nm on a Multiskan Go (Thermo scientific) plate reader using 96-well NUNC optical bottom plates. Phase separation of Y145 protein was monitored under different conditions by varying salt (NaCl) concentration (0 mM, 250 mM, 350 mM, 500 mM, 750 mM, and 1 M), temperature (4 ˚C, 15 ˚C, 25 ˚C, 37 ˚C, 50 ˚C, 65 ˚C). For temperature-dependent turbidity assays, before the measurements, the LLPS-induced solution was incubated for 5 min at respective temperatures to minimize any discrepancy due to temperature fluctuation. The sample volume used for these measurements was 150 µL and raw turbidity data are plotted without background subtraction. The formation of droplets under varying pH was estimated by recording turbidity at 350 nm on Genova Life Science spectrophotometer (ver.1.51.4). The mean and the standard error were obtained from at least three independent sets of experiments for all the measurements. For most of the experiments the Y145Stop concentration was fixed to 100 µM and salt concentration was fixed to 350 mM at pH 7.5

unless otherwise stated. To check the effect of additives such as 1,6- hexanediol and L-Arginine on LLPS of Y145Stop, the stock solution of 1,6- hexanediol (50 %) and L- Arginine (1 M) were prepared in the reaction buffer (20 mM sodium phosphate pH 7.5). The change in the turbidity was monitored by recording optical density at 350 nm using Genova Life Science spectrophotometer (ver.1.51.4). For phase separation in the presence of RNA (crude t-RNA, poly-U RNA, and yeast total RNA), 10 μM protein was used unless mentioned. RNA promoted LLPS at a protein concentration of 5 μM in 20 mM phosphate buffer with 150 mM NaCl. Phase separation was induced by adding RNA (50 ng/μL to 1500 ng/μL) to the solution (20 mM phosphate buffer, 150 mM NaCl) at room temperature. The turbidity of the solution was monitored by recording optical density at 350 nm using a Genova Life Science spectrophotometer (ver.1.51.4).

**Confocal fluorescence microscopy**

All the imaging experiments were performed on an Olympus FLUOVIEW confocal laser scanning microscope (Model No. FV10i) using a 60x oil-immersion objective (Numerical aperture: 1.35) and ZEISS LSM 710 microscope using a 63x oil-immersion objective (Numerical aperture: 1.4). To visualize droplets of Y145, 2-5 μL aliquots were withdrawn from the freshly prepared phase-separated samples and placed into a chamber made by using double-sided tape on a glass slide (Fisher Scientific 3″ x 1″ x 1 mm), which was then covered with a square coverslip. For visualizing the droplets, 1% labeled protein was doped with unlabeled protein. Y145Stop-F-5-M and Y145Stop-Alexa 488 droplets were imaged using a 473-nm laser diode (11.9 mW). Y145Stop Alexa594 droplets were imaged using an excitation source at 590 nm. All the images were processed and analyzed using ImageJ software (NIH, Bethesda, USA).

**FRAP (fluorescence recovery after photobleaching) measurements**

FRAP experiments were performed on a ZEISS LSM 710 microscope equipped with a high-resolution monochrome cooled AxioCamMRm Rev. 3 FireWire(D) camera, using a 63x oil-immersion objective (Numerical aperture 1.4). AlexaFluor488-C5- maleimide labeled protein (10 %) was used for FRAP experiments. For image acquisitions, processing, and acquiring FRAP traces, ZEN (2011) software was used. For photobleaching, a region of interest (ROI) of area = 0.61 $\mu m^2$ was bleached using a 488 nm Argon laser. The recovery of the bleached spots was recorded using ZEN Pro 2011(ZEISS) software provided with the instrument. The normalized fluorescence recovery curves were background corrected, were plotted, and fitted to estimate the half-life of recovery ($t_{1/2}$) using Origin 2018. The obtained half-life of recovery ($t_{1/2}$) and the area of the bleached ROI were then used to approximately estimate the diffusion coefficient (D) as follows: $D = \omega^2/4(t_{1/2})$.

**Circular dichroism (CD) measurements**

Far-UV CD measurements were performed on a Chirascan spectrophotometer (Applied Photophysics, UK) using a 1-mm path length quartz cuvette. Data were recorded using a final protein concentration of 10 μM protein in 20 mM sodium phosphate buffer pH 7.5. The spectra were averaged over three scans and were blank subtracted. The spectra were then processed using the inbuilt software ProData and plotted using Origin 2018.

**Estimation of saturation concentrations ($C_{sat}$)**

Centrifugation was used for estimating $C_{sat}$ as described previously.[8,9] LLPS was induced at different salt concentrations and incubated for 5 minutes at 37 ˚C to reach the equilibrium state followed by centrifugation at 37 ˚C, 16400 rpm for 30 minutes. After centrifugation, supernatants were carefully removed and pellets were dissolved in denaturant buffer (8 M Urea, 10 mM Tris, 100 mM sodium phosphate, pH 8.0). The absorbance of supernatant and pellet were recorded and the concentrations were estimated using $\varepsilon_{280} = 43{,}670\ M^{-1}cm^{-1}$.

**Steady-state fluorescence measurements**

All the fluorescence experiments were performed on a FluoroMax-4 spectrofluorometer (Horiba Jobin Yvon, NJ, USA) with a 1-mm-pathlength quartz cuvette. In all fluorescence studies, the total protein concentration used was 100 μM. For recording ANS fluorescence (10 μM of ANS) measurements, the samples were excited at 375 nm and the emission was collected in the range of 400-550 nm. For intrinsic blue fluorescence measurements, the samples were excited at 320 nm, and the emission spectra were recorded between 340 nm and 450 nm. For recording the ThT fluorescence (20 μM of ThT), the samples were excited at 440 nm, and the emission spectra were collected in the range between 460 nm and 550 nm.

**Raman Spectroscopy**

All the spectra were recorded on an inVia laser Raman microscope (Renishaw, UK). Raman spectra were recorded for Y145Stop monomer, droplets, and aggregates. The sample volume of 5 μL was deposited onto a glass slide covered with an aluminum sheet. The sample was focused using a 100x objective lens (Nikon, Japan), and a 785-nm NIR laser was used for excitation, with an exposure time of 10 s and 100% laser power. Single-droplet Raman measurements were performed by focusing the laser on individual droplets. The Rayleigh scattering was filtered by using an edge filter of 785 nm. The Raman scattering was collected and dispersed using a 1200 lines/mm diffraction grating and detected using an air-cooled CCD detector. Inbuilt Wire 3.4 software was used for data acquisition. All the data were averaged over 10 scans. Baseline correction and smoothening of the acquired spectra were performed using Wire 3.4 and the spectra were plotted using Origin.

**Atomic force microscopy (AFM) imaging**

AFM images of Y145Stop aggregates were acquired on Innova atomic force microscope (Bruker) operating in tapping mode. For sample preparation, 10 μL of the aliquots were taken from the reaction mixture and were deposited onto the freshly cleaved, Milli-Q water washed muscovite mica (Grade V-4 mica from SPI, PA). The samples were incubated for 5 minutes at room temperature and were washed twice with 100 μL of filtered Milli-Q water. The samples were further kept under a gentle stream of nitrogen gas for drying prior to AFM imaging. NanoDrive (v8.03) software was used for the data acquisition and the acquired images were processed using WSxM 5.0D 8.1 software. The height profiles were obtained from WSxM software and were plotted in Origin 2018 software.

**Seeding reactions**

For the seeding experiments, 0 h, 6 h, 12 h, and 24 h reaction mixtures (100 μM Y145Stop, 350 mM NaCl, pH 7.5; 37 ° C) were used. The samples were prepared by adding the required amounts of seeds into the fresh reaction mixture. The experiments were performed using NUNC 96-well plate, on POLARstar Omega Plate Reader Spectrophotometer (BMG LABTECH, Germany) at 25 ° C. The final concentration of ThT in the reaction mixture was 10 μM. The kinetic traces were plotted using Origin 2018.

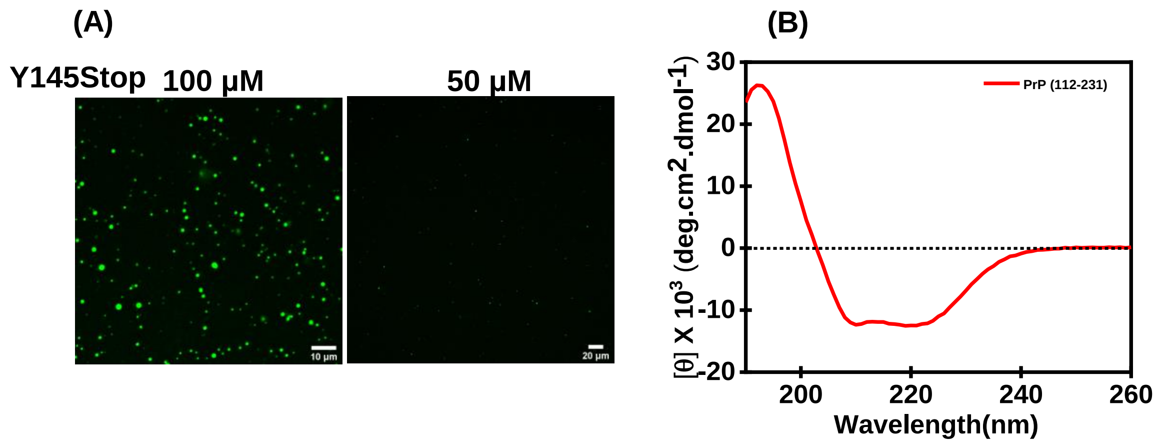


**Figure S1.** (a) Confocal fluorescence images of droplets of Alexa488 labeled Y145Stop (100 µM and 50 µM) in the presence of 10% PEG (8000 Da). (b) Far UV-CD spectrum of PrP (112-231) indicating a α-helical structure (10 µM protein, 20 mM sodium phosphate, pH 7.5).

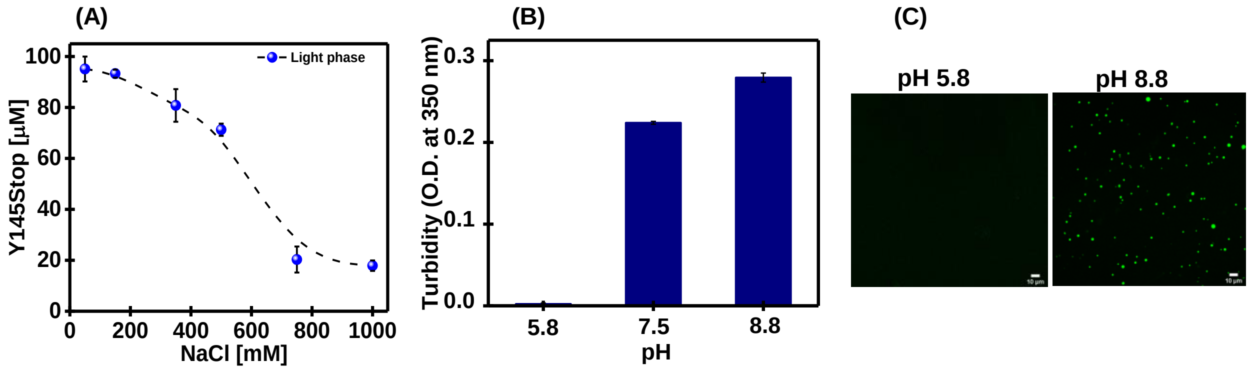


**Figure S2** (a) Concentration estimation of the (light) dispersed phase with increasing salt concentrations. (b) Turbidity plot for Y145Stop (100 µM, 20 mM sodium phosphate, 350 mM NaCl, pH 7.5) at different pH. (c) Confocal fluorescence images of Alexa488-labeled Y145Stop (100 µM, 350 mM NaCl) at pH 5.8 and pH 8.8.

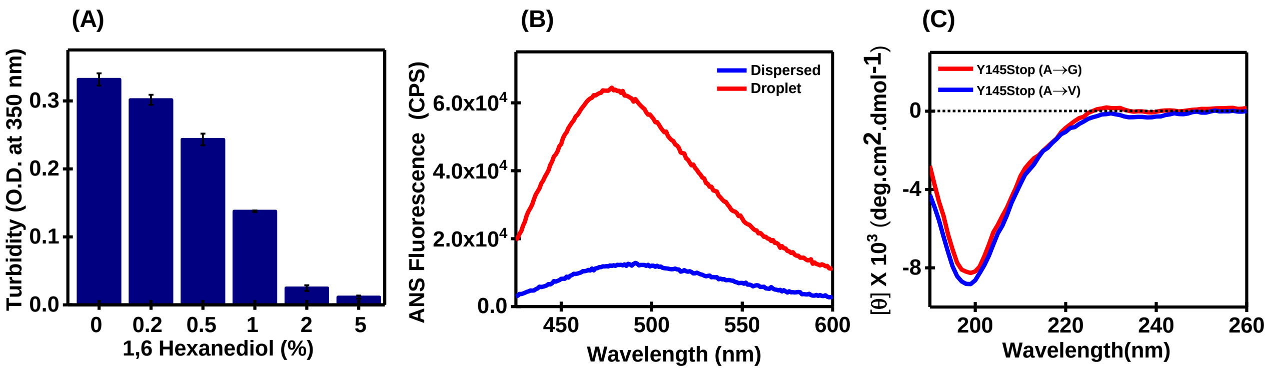


**Figure S3.** (a) Turbidity plot for Y145Stop (100 μM, 350 mM NaCl, pH 7.5) in presence of 1,6 Hexanediol. (b) ANS fluorescence for monomeric Y145Stop (black) and Y145Stop droplets (red). Inset: normalized ANS fluorescence to show the blue shift (c) Far UV-CD spectra of Y145variant, A to G and A to V (10 μM, 20 mM sodium phosphate, 350 mM NaCl, pH 7.5).

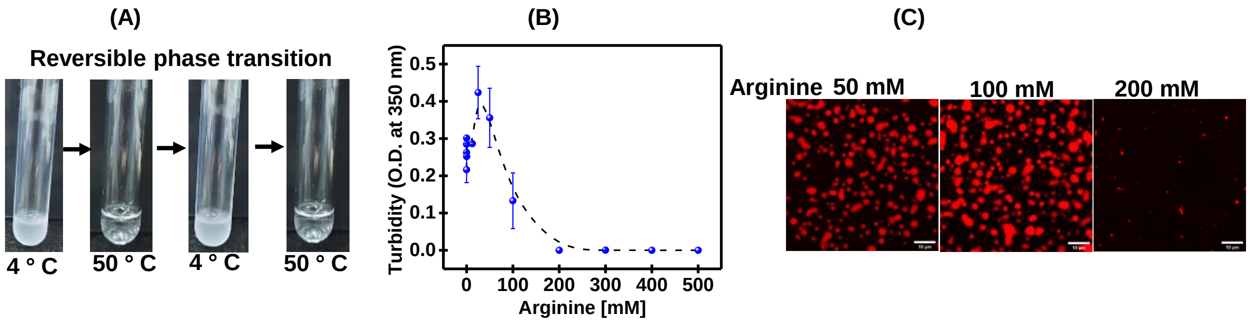


**Figure S4.** (a) Thermo-reversibility of Y145Stop droplets. (b) Turbidity plot for Y145Stop (100 μM, 20 mM sodium phosphate, 350 mM NaCl, pH 7.5) in the presence of different arginine concentrations. (c) Confocal fluorescence images for Alexa594-labeled Y145Stop at different concentrations of arginine.

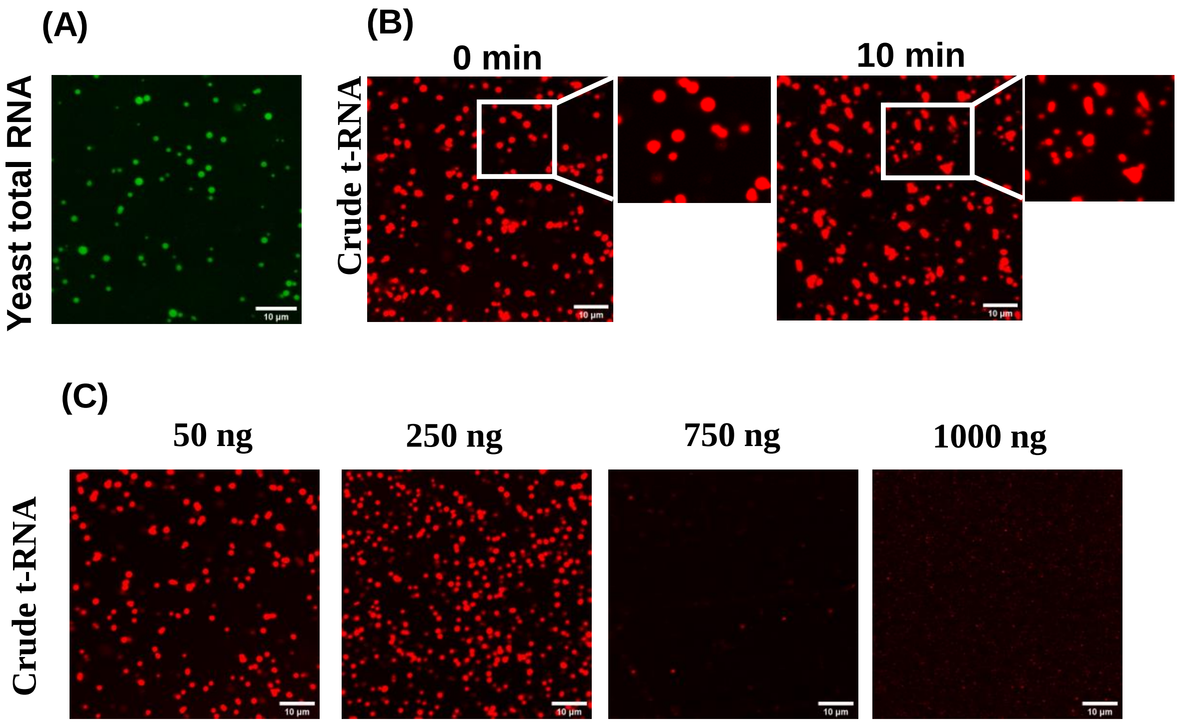


**Figure S5.** (a) Confocal fluorescence images of Alexa488-labeled Y145Stop (20 µM, 20 mM sodium phosphate, 150 mM NaCl, pH 7.5) in the presence of 100 ng yeast total RNA. (b) Confocal fluorescence images showing different morphologies of droplets formed as a course of time. (c) Confocal fluorescence images of Alexa594-labeled Y145Stop in the presence of crude t-RNA showing dissolution at high RNA concentration.

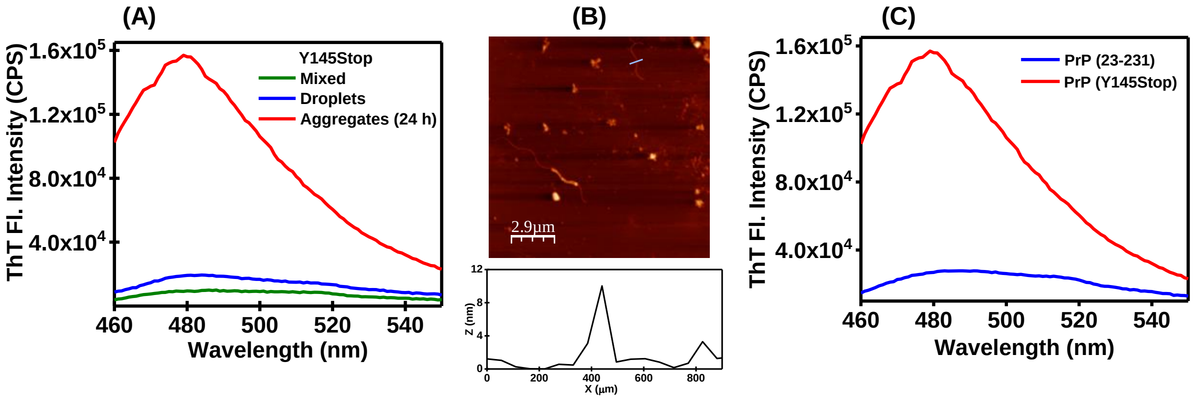


**Figure S6.** (a) ThT fluorescence intensity for Y145Stop monomeric dispersed (olive), droplet (blue), and aggregates (red). (b) AFM images of Y145Stop aggregates obtained in the presence of RNA (100 ng). (c) ThT fluorescence intensity for full-length PrP (23-231) (blue) and Y145Stop aggregates (red) obtained via aging after phase separation.

**Table S1.**

| Construct | Primer | Sequence 5'-3' |
|---|---|---|
| Y145 Stop | Forward | CGGCAGTGACTAGGAGGACCGTTAC |
| | Reverse | GTAACGGTCCTCCTAGTCACTGCCG |
| W31C Y145 | Forward | GAAGCCTGGAGGATGTAACACTGGG |
| | Reverse | CCCAGTGTTACATCCTCCAGGCTTC |
| (A113A115A117 to G113G115G117) Y145Stop | Forward | AAGCACATGGCTGGTGGTGCAGGAGCTGGGGGAGTGGTGGG |
| | Reverse | AGGCCCCCCACCACTCCCCCAGCTCCTGCACCACCAGCCATGTGC |
| (A113A115A117 to V113V115V117) Y145Stop | Forward | AGCACATGGCTGGTGTTGCAGTAGCTGGGGTAGTGGTGGGG |
| | Reverse | AAGGCCCCCCACCACTACCCCAGCTACTGCAACACCAGCCATG |
